# Membrane controlled Mechanoregulation in PIEZO1 Interactions

**DOI:** 10.64898/2026.07.16.738893

**Authors:** Sanjai Karanth, Alessandro Nicoli, Fabien Cannac, David Duria, Marina Wiesenfarth, Rita dos Santos Natividade, Julia Benthin, Dietmar Krautwurst, Andrew B. Ward, Antonella Di Pizio, Melanie Koehler

**Affiliations:** Leibniz Institute for Food Systems Biology at the Technical University of Munich, Lise-Meitner-Str.34, 85354 Freising, Germany; Department of Integrative Structural and Computational Biology, The Scripps Research Institute, La Jolla, CA 92037, USA; TUM Graduate School, TUM School of Life Sciences Weihenstephan, Technical University of Munich, Alte Akademie 8, 85354 Freising, Germany; Chemoinformatics and Protein Modelling, TUM School of Life Sciences, Technical University of Munich, Lise-Meitner-Str. 34, 85354 Freising, Germany; Atomistic Modeling Center, Munich Data Science Institute, Technical University of Munich, Garching, Germany; TUM Junior Fellow at the School of Life Sciences at the Technical University of Munich, 85354, Freising, Germany

**Keywords:** PIEZO1, SMA nanodiscs, micelles, atomic force microscopy, Yoda1 and Dooku1

## Abstract

PIEZO channels are mechanosensitive membrane proteins whose activation is governed by the surrounding lipid environment. However, the direct mechanistic contribution of native membrane composition to the molecular interactions remains unclear. In this study, a systematic comparison is made between PIEZO1 reconstituted in detergent micelles and in cell membrane–derived nanodiscs, which preserve the native lipid composition. Initial characterization employing a combination of atomic force microscopy and coarse-grained molecular dynamics simulations unveils distinct physical signatures of PIEZO1 in these two environments. Single-molecule force spectroscopy measurements demonstrate that interaction between the extracellular domain of PIEZO1 and a specific antibody exhibits unique mechanical responses strongly influenced by the surrounding membrane. In nanodiscs, PIEZO1 exhibits reversible, elastic-like behavior with preserved structural integrity and consistent adhesion forces even when modulated by Yoda1 (agonist) and Dooku1 (antagonist). Conversely, micelles induce a plastic response with altered mechanosensitivity and functional stability. Based on these findings, we propose a possible membrane-mediated force transmission pathway and quantify a simplified interaction energy landscape. Collectively, our findings offer the initial direct evidence of how the native lipid environment mechanistically governs PIEZO1 interactions, establishing native membranes as critical determinants for mechanotransduction.

## Main

Mechanotransduction is defined as the process by which mechanical forces are converted into cellular signals, thereby initiating various physiological processes. Responses generated from converting physical stimuli into electrochemical signals drive functions such as the sensation of touch, pressure, pain, nociception, proprioception, and other related somatosensations (1–3). These transduction outcomes predominantly occur through ion channels that mediate a wide range of biological effects, including vascular activity, cardiovascular function (4, 5), blood pressure regulation (6), perception of food texture in the oral cavity (7, 8), and potential therapeutic targets (9). As such, PIEZO proteins have been identified as a distinctive class of mechanoreceptors (10) with the unique property of unimodal activation triggered naturally by force (11). Structurally, PIEZO channels are large, high-molecular-weight, propeller-shaped homotrimers. An activation of these non-selective cation channels will lead to, for example, an influx of calcium ions, that trigger secondary signaling pathways. In recent years, a substantial body of research has been dedicated to the general characteristics of PIEZOs, as well as their biophysical and mechanotransductive mechanisms. This research has been thoroughly reviewed in several articles, with a few cited here (12–15). However, the fundamental question of how the cell membrane contributes to the mechanotransduction of PIEZO channels remains unanswered. More specifically, how the cell membrane influences PIEZO interactions is of particular interest. Three distinct activation mechanisms have been proposed for PIEZO activity: force-from-lipids, force-from-tether, and the membrane footprint model (11). However, with the exceptionally low packing of its transmembrane helices, its conformation is highly sensitive to its immediate physical boundary (16), indicating a strong lipid-protein interplay. While studies on PIEZOs reconstituted in lipid vesicles have confirmed the effects of membrane tension and curvature on its activity (17, 18), the empirical selection of phospholipids (maximum a ternary lipid mixture in vitro) limits the elucidation of accurate mechanotransductive responses. Understanding this concept is imperative because membrane dynamics vary with factors such as lipid packing, lipid tail length and saturation, cholesterol content, and the lipid composition of membrane rafts (19, 20). Collectively, these elements influence the physical and elastic parameters of the cell membrane, thereby directly impacting PIEZO. Hence, the objective of this study is to investigate how the physical architecture surrounding the PIEZO modulates its functionality. We achieve this by reconstituting the protein in micelles and native nanodiscs (21).

PIEZOs, also known as low-threshold mechanosensitive ion channels, have been studied to identify the minimal force required for their activity (16). However, of the different types of PIEZOs, PIEZO1 has been shown to be less rigid than PIEZO2, with mechanics favoring global membrane tension and a higher degree of conformational changes for activity (22). This finding underscores the need to clarify how PIEZOs generate and propagate forces during mechanotransduction. This knowledge gains prominence when forces are transmitted within a native membrane environment. To address this, we investigate PIEZO1 in the presence of the synthetic agonist Yoda1 and the antagonist Dooku1. Additionally, we will make theoretical estimations of the mechanical interaction energetics of PIEZO1 to gain a comprehensive understanding of the mechanotransduction mechanism.

In this study, an atomic force microscopy (AFM) approach was utilized, in which mouse PIEZO1 (mP1) was selected as the model protein and reconstituted into two distinct biomimetic systems, namely micelles and nanodiscs (**Fig. 1**). The selection of mP1 over other PIEZO variants was based on the known gating properties of mP1 and the confirmed role of membrane lipids in its modulation (23). Due to limited lipid environment, micelles provide a simplified system for studying protein-driven mechanical interactions. SMALPs (i.e. **s**tyrene **m**aleic **a**cid (SMA) **l**ipid **p**articles also called as polymer nanodiscs, here on referred to *nanodiscs*) provide the necessary native membrane composition and properties for a stable, dynamic system for interaction. These differences help us understand how lipid composition influences PIEZO1 interaction. The interactions are studied using a well-characterized antibody designed to interact with the extracellular (EC) region of the PIEZO1 Cap domain. The Cap domain plays a structural and functional role in mechanosensing and ion gating of the PIEZO1 channel (24, 25). Using AFM-based single-molecule force spectroscopy, we measured and analyzed the mechanical binding forces of interaction referred to as adhesion forces between the EC antibody and PIEZO1 protein in different biomimetic systems i.e., mP1 micelles and mP1 nanodiscs. Unless otherwise stated, ‘*binding’* refers to the interaction between the EC antibody and PIEZO1 measured by AFM-based single-molecule force spectroscopy. Additionally, we investigated the structural integrity and stability of mP1 in response to small molecule modulators (i.e., agonists and antagonists) to gain a comprehensive understanding of the PIEZO1 physiological response. Next, we suggest a force transmission pathway that explains the contribution of the PIEZO1 membrane environment to the ligand binding. To the best of our knowledge, the reconstitution of mP1 into SMA nanodiscs has not been reported previously.

**Fig. 1:**
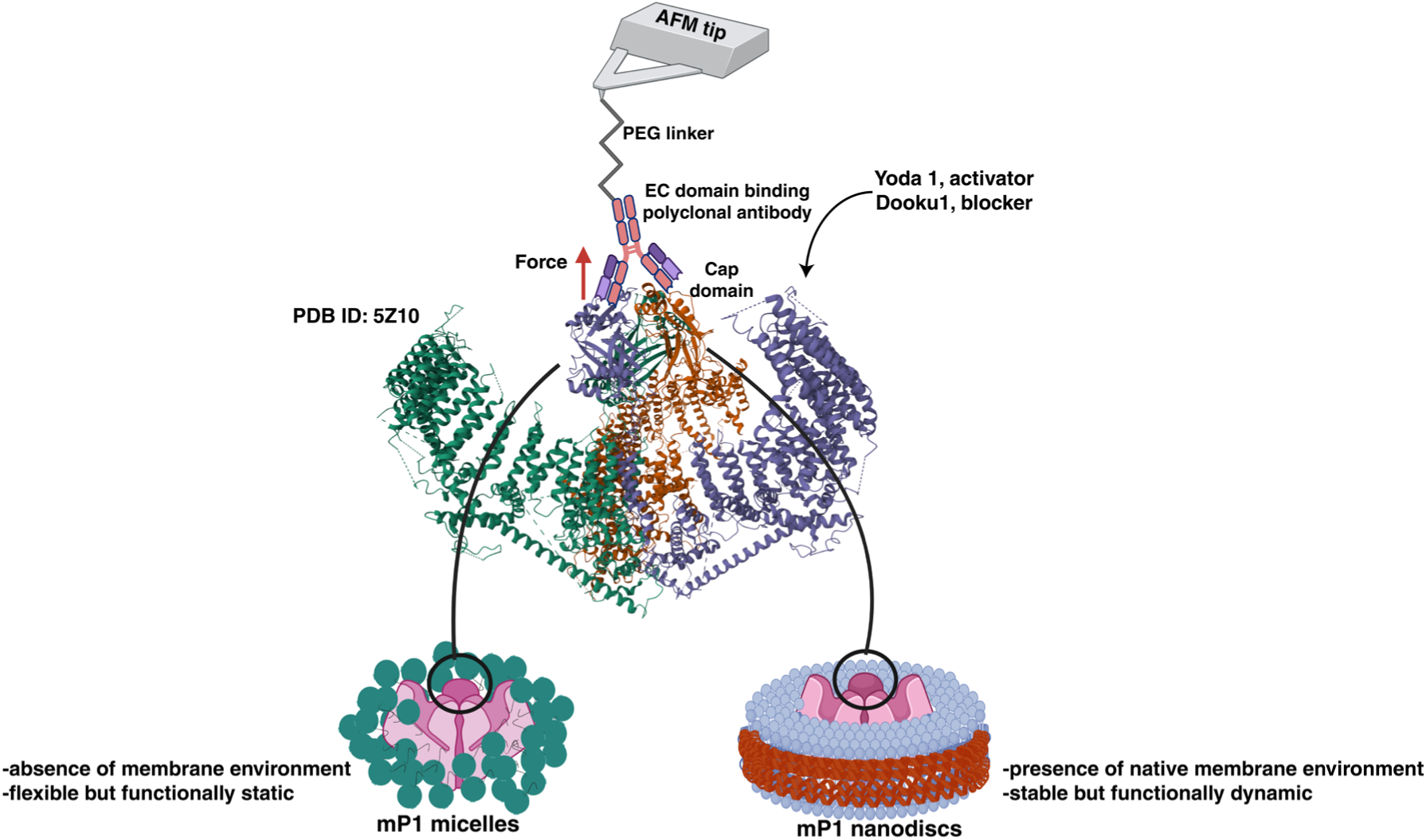
Schematic of mouse PIEZO1 (mP1) binding and molecular insights using atomic force microscopy. Reconstituted mP1 in micelles and nanodiscs exhibited distinct membrane environments that influenced the binding interaction between the EC-region polyclonal antibody and the Cap region of PIEZO1. After molecular contact, using single-molecule force spectroscopy, the binding force between the protein-antibody referred to as adhesion force was determined when the AFM tip was retracted (red arrow), generating an adhesion force between the interacting partners. Forces were measured in the absence and presence of the agonist Yoda1 and the antagonist Dooku1. In addition, the mechanical binding energetics and membrane bending energies were estimated.

Therefore, the incorporated protein was characterized in detail using coarse-grained molecular dynamics (CG-MD) simulations to provide structural context for the experimental observations. These simulations further enabled us to relate the measured mechanical responses to the intrinsic behavior of the protein within its membrane environment.

## Results

### Protein reconstituted systems show inherently distinct physical characteristics

Due to the high molecular weight of PIEZO1 and its homotrimeric nature, it was essential to verify its successful integration into micelles and nanodiscs prior to conducting binding studies. The reconstitution procedure of mP1 into micelles and nanodiscs is shown in **Fig. S1**, while initial characterization of the reconstituted systems by SDS-PAGE is presented in **Fig. S2**. As shown in **Fig. 2**, AFM imaging was used to characterize the reconstituted systems and verify the successful incorporation of mP1. AFM analysis revealed mP1 micelles (**Fig. 2a**) with a mean height of 9.0 ± 1.5 nm (**Fig. 2c**, orange box), measured from the mica surface, whereas mP1 nanodiscs (**Fig. 2b**) exhibited a mean height of 10.1 ± 1.4 nm (**Fig. 2c**, green box). While mP1 micelles were deposited directly on the mica surface, mP1 nanodiscs required mild immobilization for increased adsorption, achieved using Ni^2+^ coated mica interacting with the negatively charged SMA polymer of nanodiscs. The diameters observed for mP1 micelles are consistent with the established literature values (16). Conversely, this study is the first to report the height of the PIEZO1 protein in SMA nanodiscs. Given the mechanosensitive nature of PIEZO1, the overall protein diameter (determined as the full width at half maximum, FWHM) is an important structural parameter because membrane bending contributes directly to channel function. Additionally, unlike micelles, nanodiscs contain in-plane membrane tension, which could contribute to protein activity. The mean diameter of mP1 micelles was found to be 29.3 ± 4.9 nm, while that of mP1 nanodiscs was 25.9 ± 4.9 nm (**Fig. 2d**). The diameters observed for mP1 micelles correspond with the literature values (26). Both reconstituted systems display variation in diameter; however, in the case of nanodiscs, the contribution of the SMA polymer belt to the radial thickness should also be considered. Although the thickness of the SMA belt has been reported to be approximately 0.9 nm irrespective of nanodisc diameter (27–29), small variations may arise from differences in lipid composition. After accounting for the contribution of the SMA belt, the measured nanodisc diameter remains consistent with the incorporation of a single PIEZO1 molecule per nanodisc.

**Fig 2:**
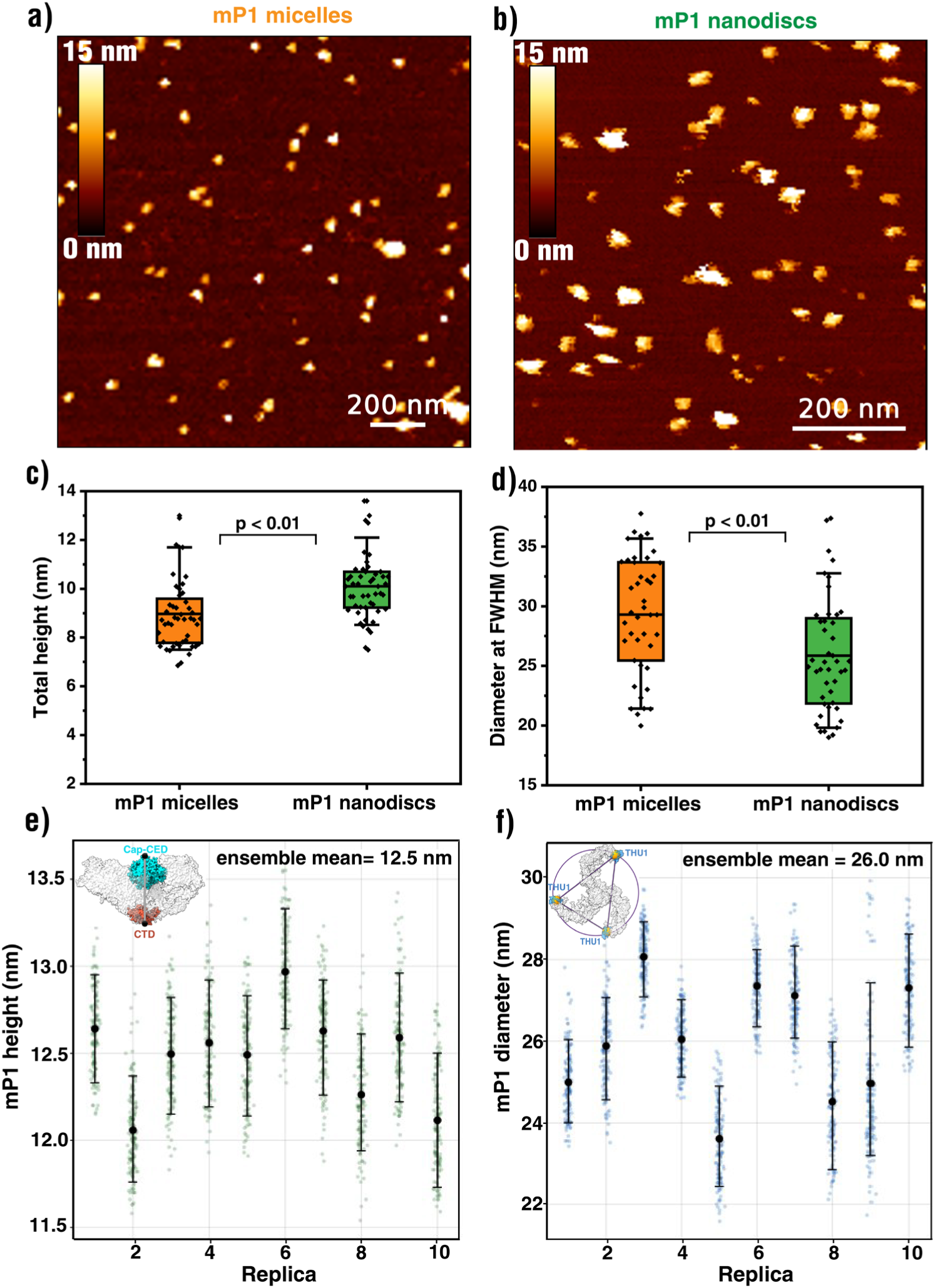
mP1 have distinct heights and diameters in different reconstituted systems. AFM liquid imaging on mica showed mP1 micelles (a) with a mean height of 9.0 ± 1.5 nm (c), while mP1 nanodiscs (b) exhibited a mean height of 10.1 ± 1.4 nm. Heights were determined relative to the mica surface, and only well-resolved particles with clearly defined contours were included in the analysis; aggregates were excluded. Diameter measurements performed at full width at half maximum (FWHM) (d) yielded mean values of 29.3 ± 4.9 nm for mP1 micelles and 25.9 ± 4.9 nm for mP1 nanodiscs. For nanodiscs, the measured diameter includes the radial contribution of the SMA polymer belt. Boxplots represent the 10th–90th percentile range, and at least 50 particles were analyzed for each condition. Along with the reported mean values, statistical analysis of height and diameter showed significant variation when two-sided t-test was performed. Protein height obtained from CG-MD simulations (e) of full-length mP1 under tensionless conditions (γ = 0 mN/m) is also reported. Transparent points represent individual simulation frames from ten independent replicas, whereas black circles and error bars denote the replica medians and corresponding 10th–90th percentiles. Inset represents the domain location with greyline representing height calculations. Triskelion diameter (f) obtained from CG-MD simulations of tensionless full-length mP1 (γ = 0 mN/m) is also shown (32). The ensemble mean across all replicas is reported in the panel for both mP1 height and triskelion diameter. Inset represents the reference positions selected for diameter estimation.

Due to the inherent membrane and SMA polymer-induced contributions to the nanodiscs, it was imperative to ascertain whether the protein dimensions in the nanodiscs are within the anticipated range and whether any conformational changes are induced. To this end, we employed CG-MD simulations to calculate the projection area of the protein, as well as its diameter and height. Although the simulations were conducted in a tensionless condition, the SMA belt has been shown to cause edge lipid radial compression (30) along with heterogeneous lipid packing-induced lateral tension. The presence of a net membrane tension (γ) in the nanodiscs can therefore be expected, depending on the relative contributions of these individual factors. Simulations across ten independent replicas yielded an ensemble mean projected protein height of 12.5 nm **(Fig. 2e**). This measurement was taken as the vertical distance between the Cap/CED and CTD regions. The diameter of the full-length tensionless mP1 was determined to be 26.0 nm (**Fig. 2f**). The calculations detailing the diameter estimation are shown in the Methods section. To understand whether nanodisc-induced tension affects the triskelion diameter, the diameter and projected area (Aproj) were predicted across different membrane tension conditions (**Fig. S3**), and linear extrapolation was carried out to observe changes in the protein diameter if a negative membrane tension prevailed within the nanodiscs. As expected, the diameter decreased with increasing compressive tension; however, the change remained below 0.5 nm even at γ = −2 mN/m. This indicates that the structure is largely preserved even under small compressive membrane stresses. Although the simulated mP1 height was higher than the experimentally obtained data for both micelles and nanodiscs, there was evidence of experimental data points within the simulated range. In contrast, the protein diameter appeared to exhibit a stronger dependence on conformational state. Compared to the estimated diameter of the free state in simulations, mP1 micelles showed a more relaxed state of the protein with an increased diameter, while the diameter of mP1 nanodiscs was similar to that in simulations. When the thickness induced by the SMA polymer belt is also factored in, the true diameter of mP1 in its native state is approximately 1.8 nm smaller than the AFM-reported value (0.9 nm belt thickness on either side). These values fall within the range reported previously by cryo-EM studies of PIEZO1 (24, 31), supporting the incorporation of a single PIEZO1 trimer per nanodisc.

### Differential PIEZO1 binding mechanics exist in micelles and nanodiscs

To investigate the influence of the membrane environment on PIEZO1 binding, AFM tips functionalized with an EC-region antibody were used to probe the samples by single-molecule force spectroscopy. Binding interactions between PIEZO1 and the antibody were monitored by recording force–distance curves during repeated approach–retraction cycles. Specific interactions were identified by a characteristic rupture event occurring after PEG-linker elongation, consistent with worm-like chain (WLC) behavior (representative example shown in **Fig. 3c**) (33, 34). The rupture force was subsequently used to determine the adhesion force. **Figures 3a** and **3b** show AFM topography images and representative adhesion maps of mP1 micelles and nanodiscs, respectively. Analysis of the antibody-PIEZO1 interaction revealed that mP1 micelles exhibited a bimodal adhesion-force distribution (**Fig. 3d**, orange). The first population was centered at 29.7 ± 1.6 pN corresponding to low adhesion, whereas the second population was centered at 145.5 ± 25.9 pN, associated with high adhesion forces. Of the two regions observed, significantly more binding events (82.2%) were recorded for high adhesion forces when 50 pN was considered the cut-off. This threshold was selected based on reported force ranges associated with supramolecular rearrangements in proteins (35). In comparison, mP1 nanodiscs revealed a single distribution with a mean adhesion force of 31.5 ± 5.6 pN (**Fig. 3d**). In mP1 nanodiscs, 79.6% of binding events occurred in the lower force region. The contribution of adhesion forces due to non-specific interactions or possible loss of the EC-antibody from the AFM tip was tested using a PEG-linker-coated AFM tip (**Fig. S4a**), as well as forces measured by the EC-antibody on empty nanodiscs made from SoyPC phospholipids (**Fig. S4b**). The measured adhesion forces in these controls ranged from 10 to 15 pN, corresponding to the noise threshold used in the analysis. The forces obtained for the nanodiscs represent molecular binding due to noncovalent interactions (36). Previously, Scheuring et al. confirmed through a high-speed AFM study that forces ranging from 25 pN to 60 pN were sufficient to induce a conformational change in the PIEZO1 protein (16). The magnitude of the forces observed in this study is consistent with previously established values.

**Fig 3:**
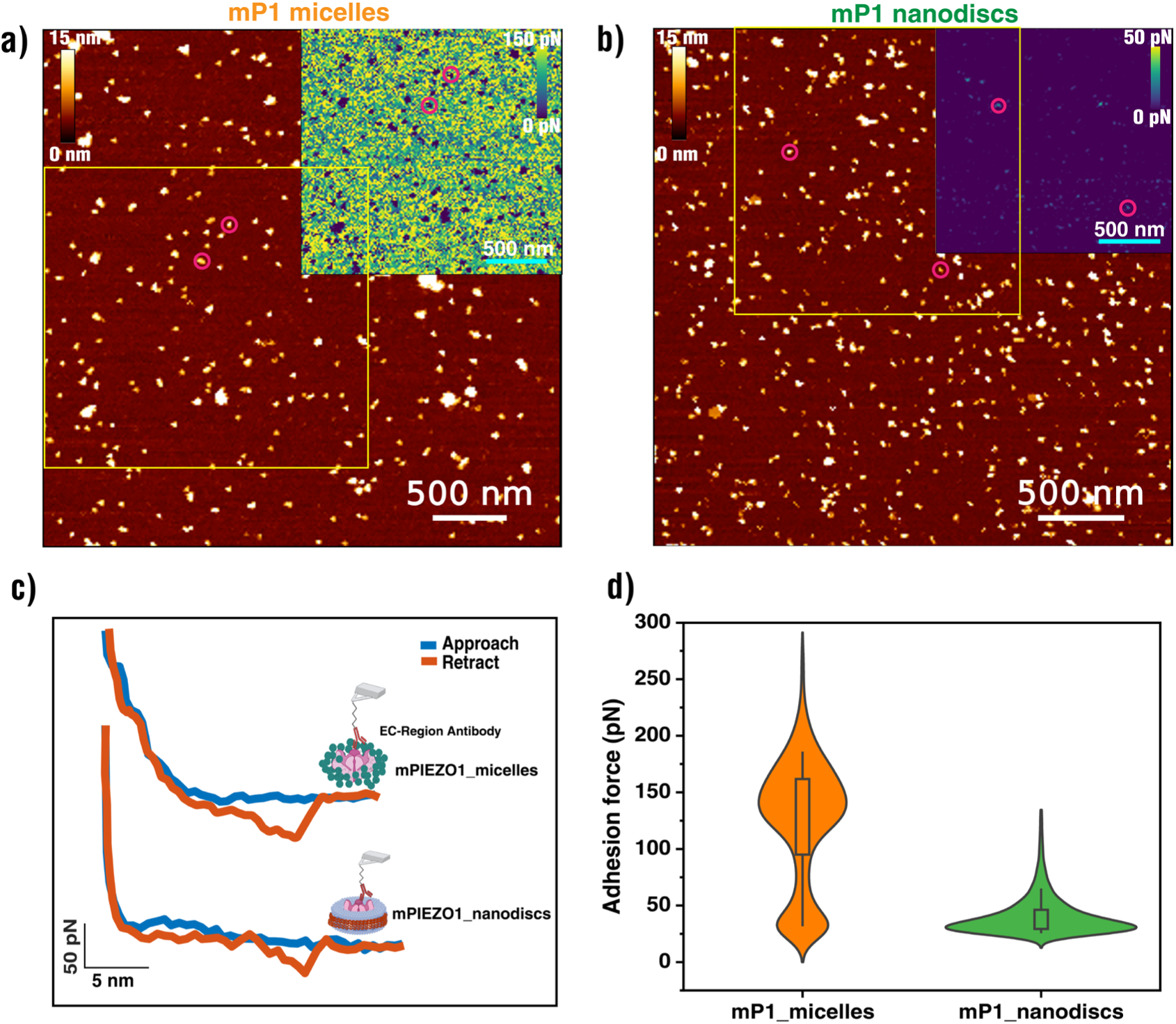
Comparative binding analysis of mP1 revealed distinct mechanical interaction profiles in micelles and nanodiscs. AFM topography images and corresponding adhesion maps acquired for mP1 micelles (a) and mP1 nanodiscs (b). Insets show representative adhesion maps from the highlighted regions, with specific binding events indicated by magenta circles. (c) Representative force–distance curves obtained during specific binding events. Adhesion forces were extracted from the retraction curves. (d) Distribution of adhesion forces measured for mP1 micelles and mP1 nanodiscs. mP1 micelles exhibited two force populations centered at 29.7 ± 1.6 pN and 145.5 ± 25.9 pN, whereas mP1 nanodiscs showed a single population centered at 31.5 ± 5.6 pN. Of the recorded binding events, 82.2% of micelle interactions occurred above 50 pN, whereas 79.6% of nanodisc interactions occurred below 50 pN. Violin plots represent kernel density estimates of the measured distributions. At least 2000 adhesion-force measurements obtained from three biological replicates, each comprising a minimum of three technical replicates, were included in the analysis. Inset Scale bar: 500 nm. Legend (mP1 micelles: 0 pN −150 pN and mP1 nanodiscs: 0 pN – 50 pN)

### Nanodiscs preserve PIEZO1 stability and ligand responses

Having established mP1 binding to the EC antibody, we next investigated how the membrane environment influences PIEZO1 activity and its mechanical binding response. To this end, we treated the mP1 micelles and nanodiscs with 10 µM Yoda1, a known PIEZO1 agonist. Computational and experimental analyses have shown that PIEZO1 possesses a binding pocket for Yoda1 in the mechanosensory arm, which is located approximately 4 nm from the central pore (37, 38). Yoda1 insertion results in the formation of a molecular wedge that effectively flattens the protein. Such flattening is expected to alter the Cap domain and consequently affect EC-antibody binding. In mP1 micelles, Yoda1 shifted the adhesion-force distribution toward lower forces, yielding a mean force of 41.6 ± 10.5 pN (**Fig. 4a**). Likewise, the mean protein height decreased to 7.6 ± 1.8 nm (**Fig. 4b**), corresponding to a reduction of approximately 14.7% relative to the untreated control. Only a small subset of particles exhibited near-complete flattening, with heights approaching 3–4 nm. In mP1 nanodiscs, Yoda1 had little effect on the adhesion forces (32.8 ± 7.3 pN) but reduced the mean height to 8.3 ± 1.8 nm (approximately 18.1% reduction compared to control), indicating activation-associated flattening while preserving the characteristic binding response. As observed in the previous system, only a limited number of complete mP1 protein flattening events were detected, with heights measured as low as 5-6 nm, which is indicative of lipid bilayer thickness.

**Fig 4:**
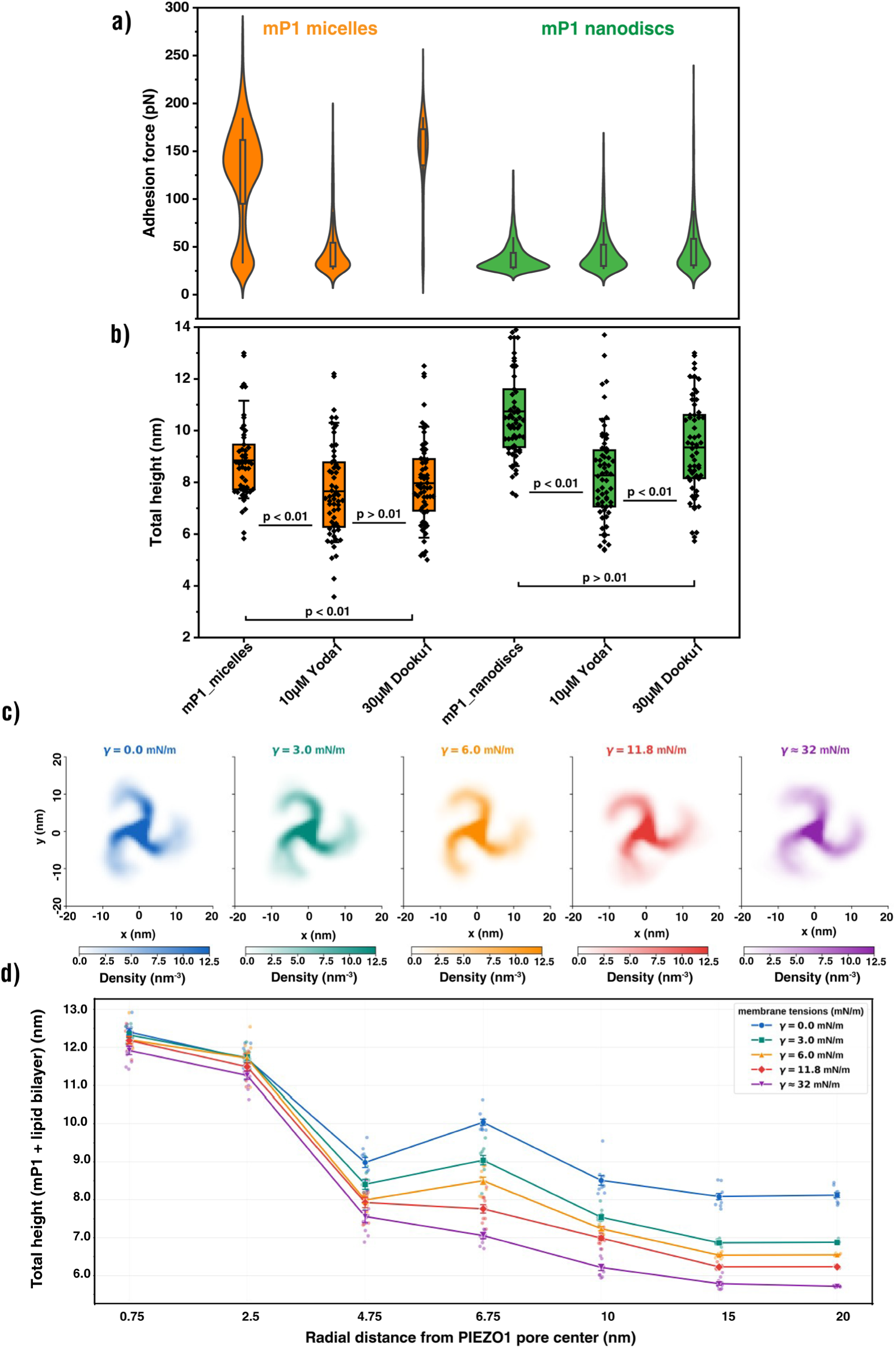
mP1 binding stability and ligand responses were preserved in nanodiscs but not in micelles. Adhesion forces measured for mP1 micelles decreased to 41.6 ± 10.5 pN following Yoda1 treatment and increased to 159.9 ± 19.2 pN after subsequent Dooku1 treatment (a). Corresponding mean heights for the treatments were 7.6 ± 1.8 nm and 7.9 ± 1.7 nm, respectively (b). For mP1 nanodiscs, adhesion forces remained largely unchanged following Yoda1 (32.8 ± 7.3 pN) and Dooku1 (34.4 ± 8.9 pN) treatment, whereas mean heights reduced to 8.3 ± 1.8 nm (Yoda1) and increased back to 9.3 ± 1.8 nm (Dooku1) (a, b). The bar graph depicts 10^th^ to 90^th^ percentile of the total data with an equal bin size maintained throughout. Horizontal line represents the mean of the bar graph. The control data denoted in 1^st^ and 4^th^ position of (a) and (b) for adhesion forces and total height has been added again from Fig 3d to benefit the readers for easy comparison. Two-dimensional projected protein density maps (c) of full-length mP1 during the simulations. Coordinates were fitted to the pore region residues 2105–2547 before projection onto the membrane plane, where x and y represent the lateral position relative to the pore center (nm). The *A_proj_* for each condition is reported in the Fig (S5, b). Height profile of mP1 with lipid bilayer (d) as a function of radial distance from the pore center, for each membrane tension condition at selected radial tranches. Data points represent mean ± s.e.m. across ten independent replicas. The complete height profiles for each tension condition are provided in Fig S6. Statistical analysis for measured heights (b) are also depicted when analyzed using One-way ANOVA test.

To investigate the observed reduction in height upon activation, CG-MD simulations were tested in two scenarios. In the first scenario, variation to the absolute protein height were analyzed across different membrane tensions. The ensemble mean height of the protein alone decreased slightly from 12.5 nm at γ = 0.0 mN/m to 12.1 nm at γ ≈ 32 mN/m (**Fig. S5a**). Dome-depth analysis revealed a progressive increase in projected area with increasing membrane tension (**Fig. S5b**), accompanied by a reduction in the protein density profile (**Fig. 4c**) highlighting the change in the shape of the protein, consistent with previously reported tension-induced flattening of PIEZO1 (32, 39–41). In the second scenario, height modulation of mP1 along with the lipid bilayer was considered. The mean total height decreased from 12.5 nm to 6 nm at the maximum membrane tension (γ ≈ 32 mN/m; **Fig. 4d**). However, at lower membrane tensions, partial flattening of the protein was detected. The height was measured as a function of radial distance from the mP1 pore center. While a full-length protein was used to interpret the conformational dynamics in the simulations, the kink at the radial distance of 6.75 nm (corresponding to the THU7 region) starts to fade away with decrease in protein height for increasing membrane tension showing structural variation to the mP1 flattening. **Fig. S6** shows the overall behavior of mP1 activation and tension-driven conformational changes for all radial positions.

Going forward, we confirmed the reversibility of protein activation by sequentially treating the samples with 30 µM Dooku1, a PIEZO1 inhibitor. Dooku1 is a small molecule that exhibits a competitive interaction with Yoda1 by replacing it. Despite targeting the same region, the depth of their binding pockets varies. Specifically, Dooku1 binds 0.3 nm above Yoda1 (42, 43). This structural variation triggers Dooku1’s antagonistic nature, thereby blocking PIEZO1 channel activation. Based on this understanding, a threefold higher concentration of Dooku1 was used. Adding the antagonist resulted in pronounced variations in adhesion forces between the two reconstituted systems. In mP1 micelles, a higher adhesion force of 159.9 ± 19.2 pN was measured compared to the Yoda1 treatment (**Fig. 4a**), indicating very strong EC-antibody binding. Furthermore, the shift in adhesion force did not correlate with reversal of PIEZO1 height, which was measured at 7.9 ± 1.7 nm (**Fig. 4b**; analogous to 7.6 ± 1.8 nm in Yoda1-activated mP1 micelles). In comparison, Dooku1 on mP1 nanodiscs exhibited adhesion forces comparable to prior measurements, with a mean of 34.4 ± 8.9 pN (**Fig. 4a**). Across all treatments, adhesion forces in nanodiscs varied by less than 5%. However, the total height increased by approximately 1.0 nm compared to the Yoda1 treatment, with a mean value of 9.3 ± 1.8 nm (**Fig. 4b**). This value is closer to the control height of the mP1 nanodiscs (10.1 ± 1.4 nm, **Fig. 2c**). The measured height also falls within the range of expected simulation data (**Fig. 4d**), as Dooku1 favors the closed state of the protein, even though it does not act directly by lowering the membrane tension. It is important to note that the contribution of the agonist or antagonist alone to the total height of the mP1 nanodiscs is negligible, as these are very small molecules with insignificant physical dimensions. The total number of recorded adhesion events for Dooku1-treated mP1 micelles was 27.9% with the EC antibody, while the mP1 nanodiscs showed 70.3% binding events.

Collectively, these results demonstrate that the native membrane environment preserved in nanodiscs supports a reversible functional modulation of PIEZO1 while maintaining a stable mechanical binding response. In contrast, micelles displayed pronounced changes in adhesion forces and impaired reversibility, indicating altered mechanosensitive behavior. To further probe the contribution of the membrane environment to PIEZO1 binding and whether force sensitivity can be influenced, mP1 nanodiscs were treated with 10 µM GsMTx4, a mechanosensitive channel inhibitor that acts primarily through membrane perturbation rather than direct interaction with the protein (44, 45). GsMTx4 treatment resulted in a broader adhesion-force distribution (**Fig. S7**) and increased the mean adhesion force to 43.1 ± 3.2 pN. Concurrently, the mean particle height decreased from 10.1 ± 1.4 nm to 9.1 ± 0.9 nm. Together, these findings demonstrate that perturbations of the surrounding membrane are sufficient to alter both the structural and mechanical response of mP1.

## Discussion

The inherent mechanosensitivity of PIEZO channels makes them attractive systems for investigating how physical forces influence their interaction with binding partners. Unlike other mechanosensitive proteins, whose activity is affected by temperature and external forces (46, 47), PIEZOs have been demonstrated to function as unimodal mechanosensors that respond only to mechanical stimuli (11). This study aimed to determine the role of native membrane composition in the mechanical response of PIEZO1 channels in two biomimetic systems. In nanodiscs, curvature and membrane-induced bending play crucial roles in PIEZO1 activity. Haselwandter et al. introduced the term "PIEZO nanodome" to describe the surrounding membrane area where, in conjunction with induced membrane bending, the protein plays a pivotal role (18). Their studies showed that increasing nanodome curvature modulates PIEZO1 activity through membrane-protein interactions. Similar membrane-coupled effects may also be expected in nanodiscs; however, unlike spherical vesicles, nanodiscs represent finite flat membrane patches confined by an SMA polymer belt. The heterogeneous native lipid composition retained within the nanodiscs, together with the large footprint of PIEZO1, likely contributes to additional curvature and local membrane stress through radial compression, thereby explaining the observed differences in height and diameter between micelles and nanodiscs. We identified distinct structural states of mP1 corresponding to the simulated free form, the detergent-associated form, and the native membrane-associated form. Based on the values obtained, mP1 micelles seem to adopt a detergent-stabilized conformation with an expanded state of mP1 compared to the more compact intrinsic structure seen in simulations. However, it is well known that detergent molecules provide a non-physiological conformation to membrane proteins, along with a hydrophobic interface that could interact with the transmembrane domains and maintain a transient state (48–50). In contrast, nanodiscs displayed a diameter comparable to the simulated intrinsic conformation, although contributions from the SMA belt must be considered when interpreting measurements. Despite these observations, unequivocal assignment of protein orientation remains challenging with the current experimental setup. The confirmed binding of the EC antibody, however, demonstrates accessibility of the extracellular domain and enabled the interaction measurements described here.

The micellar system showed higher adhesion forces during the binding of the EC antibody with the PIEZO1 protein. This behavior did not occur in the nanodisc system, which suggests distinct mechanical and structural states in the two reconstituted systems. When we estimated the binding energetics using the approximated mechanical work equation (see Supplementary **Calculation S1a**), we found that the mechanical energy required for bond rupture in mP1 micelles ranged from 86.9 k_B_T to 425.8 k_B_T (**Table 1**). In the presence of Yoda1, the estimated energy decreased to 121.6 k_B_T, whereas Dooku1 increased it to 468.2 k_B_T. Based on the estimated energetic landscape, PIEZO1 in micelles appears to populate at least two mechanically distinct states, reflected by the bimodal adhesion-force distribution. However, no such shift in energetics was observed in nanodiscs. The transitional behavior observed in micelles in the presence of an agonist and a antagonist is consistent with their well-defined, state-dependent binding (32, 42). Yoda1 reduces the intrinsic curvature of PIEZO1 and lowers the energetic cost of flattening, an effect that is partially reversed by Dooku1. Nevertheless, the affirmation is limited by the functional aspect of the protein, as evidenced by the inability to reverse the protein height after adding Dooku1.

**Table 1:** Estimated mechanical binding energetics based on mP1-EC antibody interaction. Binding energetics were estimated from the mean adhesion forces measured in Fig. 3d and Fig. 4a and an average tip-sample separation of 12 nm. Details of the calculations are provided in Calculation S1a (binding energetics) and Calculation S1b (membrane bending energy based on the Helfrich elasticity model). * Membrane bending energy was estimated independently of treatment because the overall nanodisc dimensions are constrained by the SMA polymer belt. Therefore, bending energies are reported for the two limiting possible geometries in this study, namely elliptical and circular nanodiscs. Calculations were performed at 298 K and for k_B_T conversion ^§^. Two values are reported for mP1 micelles because the control condition exhibited two distinct adhesion-force populations. Energetic estimates shown here are intended to provide comparative values between conditions rather than statistical confidence intervals.

| Reconstituted system | Estimated binding energetics (in k<sub>B</sub>T) |  |  |
| --- | --- | --- | --- |
| | Control | 10 μM Yoda1 | 30 μM Dooku1 |
| mP1 micelles | 86.9 and 425.8<sup>§</sup> | 121.6 | 468.2 |
| mP1 nanodiscs | 107.4 | 111.9 | 117.4 |
| Membrane bending energy (in nanodiscs) | 48.1 (elliptical shape) – 11.9 (circular)* |  |  |

On the other hand, the mechanical profile of mP1 nanodiscs showed a significant degree of independence from the treatment method. Estimated binding energies of mP1-antibody in nanodiscs changed only marginally, increasing from 107.4 k_B_T in the control condition to 111.9 k_B_T and 117.4 k_B_T following Yoda1 and Dooku1 treatment, respectively.

Unlike micelles, the energetics of nanodiscs are also attributed by the bending energy of the membrane and the radial compression of the SMA belt. The bending rigidity of a free-standing lipid bilayer has been estimated to be 20 k_B_T using the Helfrich model of membrane elasticity (51, 52). This estimation applies to simple lipid mixtures. However, nanodiscs have an anisotropic composition, which increases the bending rigidity to 35 k_B_T for large diameters (53). Accordingly, the total membrane bending energy of an elliptical nanodisc can be estimated to be 48.1 k_B_T (see Supplementary **Calculation S1b**) showing that nearly half of the total mechanical work done by the protein comes from curvature- or deformation-induced interaction effects alone. The presented finding underscores the substantial contribution of the membrane to the mechanical response of PIEZO1 and highlights the intimate coupling between the protein and its surrounding lipid environment. This interpretation is further supported by the GsMTx4 experiments, where perturbation of the membrane environment produced measurable changes in adhesion forces. Together, these observations suggest that small-molecule modulators induce subtle changes in native-state PIEZO1 interactions without causing large-scale alterations in the adhesion-force landscape. Such behavior is consistent with a mechanically controlled system that preserves PIEZO1 dynamics and conformational responses compared to micelles. Furthermore, the observed stability of the developed nanodiscs system while maintaining functional modulation could have direct implications in pharmacodynamic studies, as it may assist and possibly overcome the limitations of canonical ion channel assays (e.g., fluorescence related anomalies in calcium signaling readouts).

## Conclusion

Overall, the study highlights fundamental differences in the environment-dependent mechanical coupling of PIEZO1 and their consequences during binding. **Fig. 5** summarizes the observed protein interaction response. In a system lacking a membrane environment, such as in micelles, PIEZO1 exhibits rigid mechanical coupling. In such system, interaction forces and their transmission are expected to be dominated by the intrinsic mechanical properties of the protein and its surrounding detergent shell, with only limited dissipation into the micelles. In contrast, in a nanodisc system, PIEZO1 is mechanically coupled to the membrane geometry. This coupling enables the creation of a "buffered" state, in which membrane undulations redirect and redistribute force propagation rather than acting directly on the protein interface, as reflected by the lower adhesion forces. This observation is consistent with the membrane dome model (17), in which the physical load is distributed across the membrane-protein assembly, thereby preserving the native mechanical behavior of PIEZO1.

**Fig 5:**
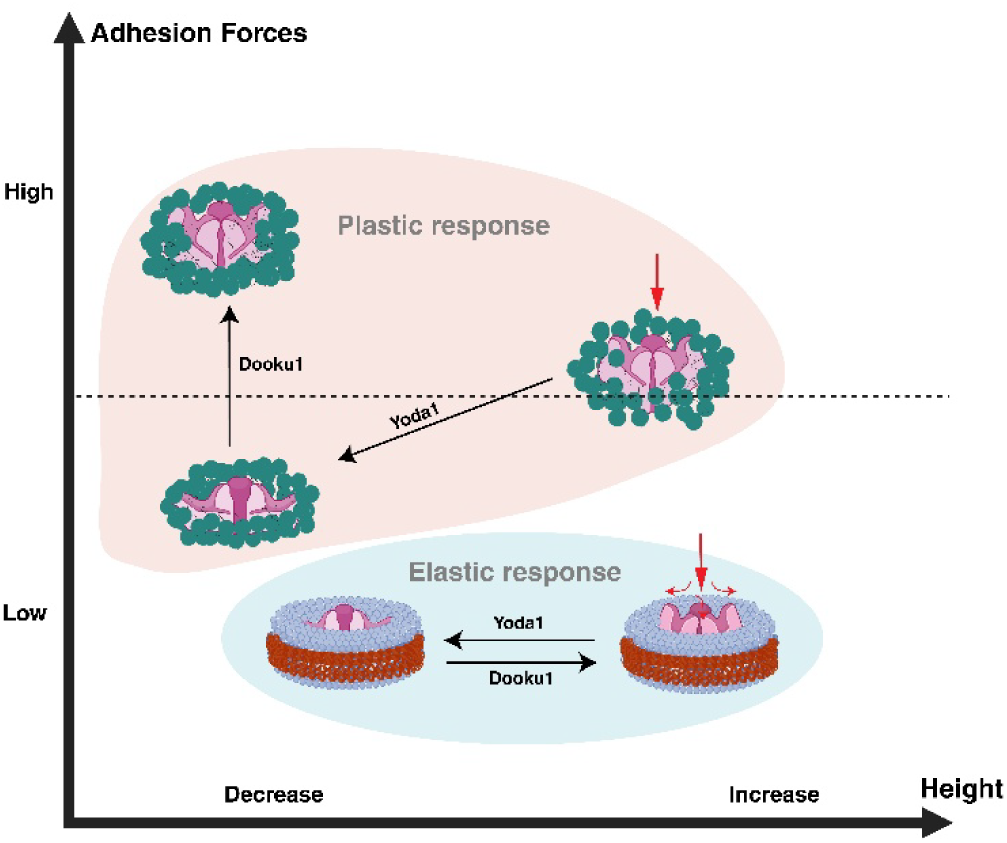
A proposed interaction model on the role of membrane environment in PIEZO1 binding mechanics. In micelles, PIEZO1 is surrounded by a detergent shell that provides limited membrane-mediated mechanical coupling. As a result, changes induced by modulators are reflected in large shifts in adhesion forces and altered structural behavior, consistent with a plastic mechanical response. In contrast, nanodiscs provide a membrane environment that remains mechanically coupled to the protein. This membrane coupling is proposed to redistribute local mechanical perturbations and maintain a stable binding-force regime despite activation- and inhibition-dependent conformational changes. The resulting behavior is consistent with an elastic mechanical response, in which membrane-associated force transmission contributes to the preservation of native-like PIEZO1 function.

The presence of modulators such as Yoda1 and Dooku1 alters the internal mechanics of the protein directly within micelles. This alteration likely changes the protein–micelle contact or packing geometry, which influences the overall spatial distribution of the protein. Depending on the modulator, the mechanical coupling is either reduced (as with Yoda1, which lowers adhesion forces) or increased (as with Dooku1, which increases adhesion forces), making the protein–micelle interaction more rigid. The resulting changes in protein conformation and detergent coupling may disrupt the native mechanical context of the ion channel, which explains the drastic shifts in adhesion forces and the ‘plastic’ response observed in micelles. However, in nanodiscs, these modulators act on an existing "buffered" system. Consequently, the altered mechanics cannot overcome the protein–membrane coupling geometry to induce an out-of-plane deformability, despite the fact that binding of Yoda1 or Dooku1 to PIEZO1 causes conformational changes. In other words, the membrane geometry acts as a mechanical buffer that preserves the overall dynamic behavior of PIEZO1 despite local perturbations, consistent with the low and largely unchanged adhesion forces measured in nanodiscs. This behavior suggests that, under native conditions, PIEZO1 force transmission operates with an ‘elastic’ response, where tuning of the gating equilibrium occurs rather than a complete reorganization of its mechanical pathway. This outcome is consistent with the theoretical understanding that mechanosensitive channels exploit membrane mechanics for efficient force transmission.

Taken together, our findings demonstrate that the membrane environment is not merely a passive scaffold for PIEZO1 but an active determinant in its mechanical binding behavior. Native membrane-derived nanodiscs preserve reversible functional modulation while maintaining a stable mechanical binding profile, revealing a previously unrecognized role of membrane architecture in regulating PIEZO1 mechanotransduction.

## Materials and Methods

### Transfection of HEK293 cells for mPIEZO1 expression

Transient transfection to purify the mP1 protein for reconstitution into micelles was adapted from Saotome et al (24). HEK293F cells with a density of ∼1.5 million/mL (purchased and authenticated from ATCC, tested negative for mycoplasma) were grown at 37° C, 5.0% CO_2_ and transfected with a pcDNA3.1-IRES-GFP plasmid containing mouse Piezo1 (Uniprot KB accession number: E2JF22) fused to PreScission protease cleavage site (LEVLFQGP) and GST coding sequence. 300 μg of DNA and 900 μg of polyethylenimine were used per liter of transfected HEK cells. 4 mM sodium butyrate was added to the cells 12–24 hours after transfection. The cells were then harvested and resuspended in buffer after 48 hours for reconstitution in mP1 micelles.

To purify and reconstitute the mP1 protein into nanodiscs, a large-scale adaptation of the transfection procedure with increased yield was needed. Accordingly, the transfection protocol was modified by seeding a total of ∼5 million HEK293T cells in a large TC175 culture flask (Greiner Bio-one) at 37° C, 5% CO2, and allowed to reach 70-90% confluency. On the day of transfection, a tube containing 50 ng of plasmid DNA in TE buffer was mixed with 2.4 mL of Opti-MEM (Gibco) having 1 % Penicillin-Streptomycin (Gibco) (P/S) which was preheated to 37°C. In another tube, 172.5 µl of transfection reagent, Lipofectamine 2000 (ThermoScientific), was mixed with 2.4 mL Opti-MEM. Both tubes were gently combined and incubated for 10 min at RT. The medium of the HEK293T cells was replaced with fresh DMEM + GlutaMAX (Gibco) (10% FBS, 1% P/S), and the DNA-Lipofectamine mixture was added dropwise to the cells with gentle swirling and incubated at 37°C, 5% CO₂ for 24 h. The culture media was exchanged with fresh media, and sodium butyrate was added to the flask, for a final concentration of 8 mM, and incubated overnight. The cells were then collected the next day by aspirating the media out and washing the cells once with PBS buffer, then collecting them by resuspension with PBS buffer, and pelleting them by centrifugation at 215g for 3 min. After removing the supernatant, the cell pellet was immediately frozen at −20°C. On achieving a total number of around 150 million cells from 5 - 6 different flasks, the cell pellet was processed for reconstitution of mP1 into SMA nanodiscs.

### Reconstitution of mPIEZO1 into micelles

The mP1 reconstitution was carried out as outlined in Saotome et al (24). Briefly, the transfected cells were collected and resuspended in buffer containing 25 mM HEPES, pH 8.0, 300 mM NaCl, 1% CHAPS, 1% C12E9, 2 mM DTT, and protease inhibitor cocktail (Roche). The cell lysate was stirred for 1 h at 4°C, and the supernatant collected after centrifugation was incubated with Glutathione Sepharose 4B resin (GE Healthcare) for 3 hours. The resin was washed extensively (50 CV of GST resin bed volume) with buffer containing 25 mM HEPES, pH 8.0, 150 mM NaCl, 100 μM glyco-diosgenin (GDN), and 2mM DTT. The sheer volume involved in the washing steps effectively exchanged C12E9 detergent with GDN. Post affinity-tag purification, PreScission protease (Cytiva) was added to a ∼30% slurry of resin in GDN containing buffer at a ratio of ∼100 μg per 1 mL resin and incubated at 4°C overnight. Tag-free mP1 micelles were then collected from the flowthrough, concentrated, and stored at −80°C. Due to the extensive washing steps involved and further sample dilution required during the AFM measurements, size exclusion chromatography was avoided to prevent additional sample loss. Tris-HCl buffer was used from here on for dilutions as it was the stable buffer for both reconstituted systems.

### Reconstitution of mPIEZO1 into SMA nanodiscs

For reconstituting mP1 into SMA nanodiscs, the transfected cell pellet was washed once with PBS buffer (VWR) by centrifugation at 3000 × g for 5 min. After discarding the supernatant, the cell pellet was resuspended in an ice-cold hypotonic solution (1:3 pellet volume) consisting of 10 mM Tris-HCl (Carl Roth), pH 7.4, 5 mM DTT (Thermo Scientific), and 1× protease inhibitor cocktail (Thermo Scientific). The suspension was incubated on ice for 15 min and subsequently extruded through a 26-gauge syringe needle (10–15 passes). Cell debris was removed by centrifugation at 9000 × g for 30 min at 4°C. The resulting supernatant was subjected to ultracentrifugation at 125000 × g for 60 min at 4°C to pellet the membrane fraction. The membrane pellet was resuspended in 500 µL of buffer A (25 mM Tris, 150 mM NaCl, pH 7.4; Carl Roth). A 1% SMA polymer solution (SMALP-200, Cube Biotech) was added to the membrane suspension, and the mixture was incubated for 12 h at 21°C with gentle mixing (300 rpm) to allow nanodisc formation. Excess SMA polymer and free lipid material were removed by size-exclusion chromatography (SEC) using a Superdex 200 Increase 10/300 GL column equilibrated with buffer A. The collected fraction was concentrated and subjected to GST affinity purification using a washed Glutathione Sepharose 4B resin slurry (30% slurry in 50 mM Tris-HCl, 140 mM NaCl, pH 7.4; Serva). The nanodiscs were incubated with the resin for 4 h at 4°C with gentle mixing (300 rpm), followed by centrifugation at 1500 × g for 5 min. Bound mP1 nanodiscs were subsequently eluted using freshly prepared elution buffer (50 mM Tris-HCl, 140 mM NaCl, 10 mM reduced glutathione, pH 8.0) for 15 min at room temperature. Following centrifugation at 1500 × g for 5 min, the eluted mP1 nanodiscs were recovered in the supernatant. PreScission protease was added, and the sample was incubated overnight at 4°C without mixing. A final SEC purification step was performed after concentrating the sample to 100 µL using a 30 kDa MWCO filter (Sartorius) and loading it onto a Superdex 200 Increase 10/300 GL column. Peak fractions were collected at a flow rate of 0.5 mL/min and stored at 4°C. Purified mP1 nanodiscs were used for AFM measurements within one week of reconstitution.

### AFM measurements

#### Functionalization of AFM tips with the EC-region antibody

AFM tips were functionalized with a polyclonal antibody directed against the extracellular (EC) domain of mP1 (15939-1-AP, Proteintech). This antibody has been extensively characterized for its interaction with PIEZO1 (54–56). Antibody immobilization was performed according to a previously established protocol (57). Briefly, BL-AC40TS silicon nitride cantilevers (Olympus Corporation) were cleaned in chloroform (Carl Roth) for 5 min, dried under N₂, and subsequently treated with UV-ozone (NanoBioAnalytics) for 10 min. The cleaned tips were amino-functionalized with 3-aminopropyltriethoxysilane (APTES, Sigma) in the gas phase using an argon-filled desiccator containing 30 μL APTES and 10 μL triethylamine (Sigma) in separate reservoirs. Cantilevers were positioned adjacent to the reagents and incubated for 2 h. The reagent reservoirs were then removed, and the cantilevers were left in the argon atmosphere for an additional 2 days to cure the APTES coating. PEG linker molecules were attached by dissolving 3.3 mg aldehyde-PEG-NHS (BroadPharm, Mw ≈ 1375.6 g/mol) in 0.5 mL chloroform and mixing with 30 μL triethylamine. The cantilevers were transferred to a reaction chamber and incubated for 2 h, followed by washing with chloroform (3 × 5 min). After drying under N₂, the cantilevers were arranged in a circular pattern in a Parafilm®-covered Petri dish with the tips facing inward. A freshly prepared solution of 1 M sodium cyanoborohydride (Sigma) in 20 mM NaOH (Carl Roth) was used for antibody coupling. Subsequently, 100 μL of a 0.1 mg/mL antibody solution was applied to the cantilevers, followed by the addition of 2 μL of 1 M sodium cyanoborohydride. The reaction mixture was incubated for 1 h. Thereafter, 5 μL of 1 M ethanolamine (Sigma, pH 8.0) was added and incubated for 10 min to quench the reaction by blocking residual aldehyde and NHS groups. The cantilevers were subsequently washed with HEPES buffer (25 mM HEPES, 150 mM NaCl, pH 7.4; 3 × 5 min) and stored at 4°C. Functionalized cantilevers were used within one week of preparation.

#### Single molecule force spectroscopy (SMFS)

AFM-based SMFS experiments (BioAFM JPK Nanowizard V, Bruker) were performed on mP1 micelles and mP1 nanodiscs with EC-antibody functionalized tips at RT (∼23-25°C). Unless otherwise stated, all experiments were conducted in buffer containing 25 mM HEPES and 150 mM NaCl at pH 7.4. For SMFS measurements, mP1 micelles (∼1–5 µg/mL) were deposited onto freshly cleaved mica and incubated for 10 min at RT. The surface was subsequently washed 10–15 times with buffer to remove unbound micelles while maintaining a sparse surface distribution of mP1. In contrast, mP1 nanodiscs exhibited limited adsorption to mica and therefore required immobilization. For this purpose, Ni²⁺-modified mica was prepared by incubating freshly cleaved mica with 100 µL of 1 mM NiCl₂ (Sigma) prepared in Milli-Q water for 5 min. After washing the surface three times with Milli-Q water, mP1 nanodiscs were added and incubated for 25 min. Unbound nanodiscs were removed by washing the surface 5–7 times with buffer. Ni²⁺ ions served only for immobilization along with minimizing substrate-induced effects on the physical and mechanical measurements. SMFS measurements were performed in QI mode, in which force– distance curves are acquired at each image pixel while the functionalized tip sequentially probes the sample surface for interactions with PIEZO1. This approach generates topography images and adhesion maps simultaneously. The resulting force data, corresponding to individual approach–retraction cycles, were compiled into adhesion maps with a resolution of 256 × 256 pixels. Relatively large scan areas were selected to capture the spatial distribution of mP1 particles and maximize the number of detected binding events despite the long acquisition time associated with adhesion mapping. The spring constant of each functionalized cantilever was calibrated prior to every experiment using the thermal noise method. BL-AC40TS cantilevers (Olympus, Japan) were operated with a setpoint force of 300 pN and spring constants ranging from 0.08 to 0.10 N/m. Because PIEZO1 functions as a force transducer, a previously validated setpoint was employed (54).

To investigate the activation effect of mP1, 10 µM Yoda1(58) (Sigma) was freshly prepared in buffer from a 1 mM stock solution in DMSO stored at −20°C. The Yoda1 solution was introduced by buffer exchange and incubated with mP1 micelles or nanodiscs for 15 min at RT. Subsequently, the solution was exchanged twice with fresh buffer prior to SMFS measurements. For blocking studies, 30 µM Dooku-1 (59) (Sigma) was freshly prepared in buffer from a 1 mM stock solution in DMSO stored at −20°C and added to the Yoda1-treated samples. Following incubation for 15 min at RT, the solution was exchanged twice with fresh buffer before continuing the SMFS measurements. In GsMTx4 experiment, the mP1 nanodiscs were treated with 10 µM GsMTx4 for 15 min followed by SMFS for adhesion force.

In addition to the internal controls provided by parallel topography imaging and adhesion mapping, two external controls were performed: (i) measurements using AFM tips functionalized with PEG linker only and (ii) measurements using EC-antibody-functionalized tips on empty nanodiscs.

#### Data analysis

Raw AFM image processing and force–distance (FD) curve analysis was performed using SPM Data Processing software (v8.0.85, Bruker). For the physical characterization of mP1 micelles and nanodiscs, only isolated particles exhibiting a well-defined morphology and flat adsorption onto the substrate were selected for height and diameter analysis. Total particle height was measured relative to the mica surface. The contour of each particle was identified, and the lateral diameter was determined using the full width at half maximum (FWHM) of the corresponding height profile. Only particles with clearly defined contours were included in the analysis. Following this p value analysis was carried out using two-sided t-test with significance level at 0.01. For FD curve analysis, binding events corresponding to interactions between mPIEZO1 and the EC antibody were identified from individual adhesion maps. Adhesion forces were determined from the minimum force observed in the retraction segment of each force–distance curve. To exclude contributions from instrumental noise and environmental artifacts, a force threshold between 10 and 20 pN was applied. Consequently, only binding events exhibiting adhesion forces greater than 20 pN were considered for further analysis and exported to OriginPro 2021b (OriginLab) for statistical evaluation. Data distributions were visualized using violin-box plots with uniform binning and analyzed by kernel density estimation to identify distinct force populations. Local mean values within the distributions were determined using nonlinear multi-peak Gaussian fitting. The corresponding height analysis for the measured binding events were performed as before and p value analysis was performed using One-way ANOVA with significance at 0.01.

### Computational investigation of mP1

Coarse-grained (CG) molecular dynamics (MD) simulations of mouse PIEZO1 (mP1) embedded in a heterogeneous lipid bilayer were retrieved from the public repository at https://doi.org/10.17617/3.USULVG. Each system contained ten independent replicas for each membrane tension condition. Each replica produced a 4 μs trajectory, with frames saved every 20 ns, yielding 201 frames per replica (32). Raw trajectories were post-processed using GROMACS (trjconv) (60), followed by atom reordering to ensure sequential residue numbering across the three chains of the PIEZO1 homotrimer (Uniprot accession code: E2JF22). All trajectory analyses were performed using a custom Python (3.12) pipeline built on MDanalysis (v 2.2.0) (61), with data visualization using Matplotlib (v3.10.9) and Seaborn (v0.13.2).

The dome projected area (*A_proj_*) was computed for each frame as 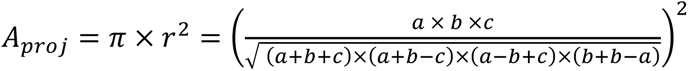, where *r* is the radius of the projected area, using the center of geometry (COG) of the outermost resolved repeat of each monomer (THU1, residues 13–138) as the three vertices, where a, b, and c are the pairwise distances between the three THU1 COGs(40, 41). The corresponding triskelion diameter was defined as *D* = 2 × *r*. Protein height was computed per frame as the z-distance between the lowest backbone bead of the Cap/CED domain (residues 2213–2457) and the highest backbone bead of the CTD (residues 2486–2547), both measured relative to the pore center of mass.

Similar to the approach described by Chong et al (39) for mP1, the aligned trajectories were projected onto the membrane plane to generate averaged two-dimensional protein density maps. Backbone (BB) bead positions were projected following rotational alignment to the pore region (residues 2105–2547) and accumulated on a symmetric 75 × 75 Cartesian XY grid centered on the pore center of mass. Densities were normalized by bin area and total number of frames to obtain maps in units of nm^-3^. Density maps were averaged across all frames and replicas and Gaussian-smoothed for visualization. The radial height profile of the mP1–membrane complex was computed as a function of radial distance from the pore center. Protein and lipid beads were assigned to concentric annular bins of 0.5 nm width extending to a maximum radius of 25 nm. For each annulus, the local height was defined as the full z-span (maximum minus minimum z-coordinate) of all protein and lipid beads within the bin. Profiles were averaged over all post-equilibration frames and are reported as ensemble mean ± s.e.m. across the ten replicas.

## Data availability

All the data presented in this study are included in the manuscript and supplementary information. Any additional information needed can be directed to the corresponding authors.

## Author Contributions

S.K: Project conceptualization, developed and characterized nanodiscs, AFM experiments, Data analysis, Project coordination, Manuscript original draft, review and editing.

A.N: Data analysis of MD simulations, Visualization, Manuscript review and editing.

F.C: Developed and characterized micelles systems, Manuscript review and editing.

D.D, M.W, D.K, R.S.N and J.B: Cell culture and Transfection, Manuscript review and editing.

A.B.W and A.D.P: Manuscript review and editing.

M.K: Project conceptualization and coordination, Funding, Manuscript review and editing.

## Ethics declarations

The authors declare that they have no known competing financial interests or personal relationships that could have appeared to influence the work reported in this paper.

## Supplementary information

Supporting information to this manuscript include Fig S1: showing the schematic overview on the mP1 reconstitution procedure, Fig S2: Characterization of the reconstituted mP1, Fig S3: Estimation of mP1 diameter and projected area under different membrane tensions from CG-MD simulations, Fig S4: Control SMFS measurements for segregating non-specific interactions, Fig S5: Height and area analysis of full mP1 under different membrane tensions as observed through simulations, Fig S6: Radial height profile of mP1 embedded in the lipid bilayer under different membrane tensions, Fig S7: Effect of GsMTx4 treatment on mP1 nanodiscs and Calculation S1: Estimation of Binding energetics and membrane bending energy.

