## Supplementary File for "Membrane controlled Mechanoregulation in PIEZO1 Interactions"

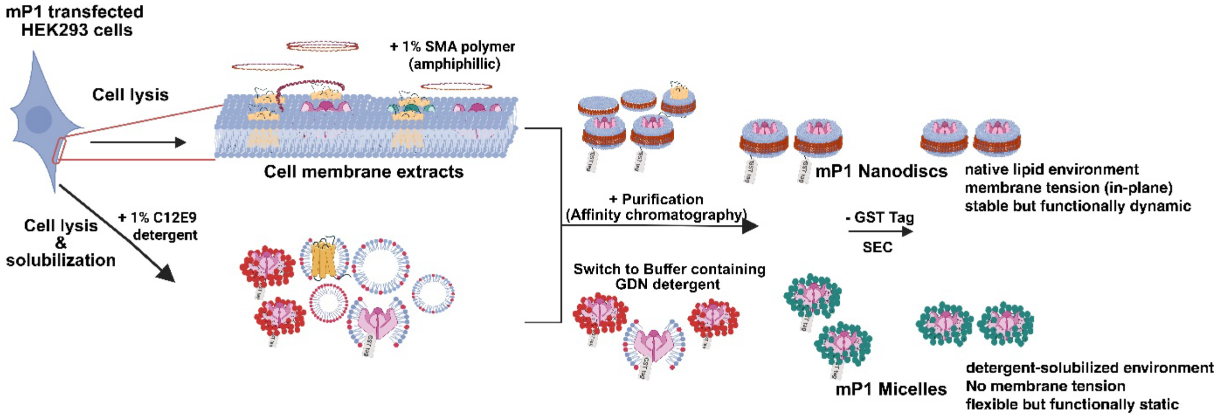


**Fig S1: Schematic overview of mouse PIEZO1 (mP1) reconstitution into micelles and SMA nanodiscs.** mP1-transfected HEK293 cells were lysed and processed using two different reconstitution strategies. For micelle preparation, membrane proteins were solubilized using C12E9 detergent followed by GST-affinity purification. For nanodisc preparation, membrane extracts were treated with 1% styrene–maleic acid (SMA) polymer to generate SMA lipid particles (SMALPs), followed by GST-affinity purification and size-exclusion chromatography (SEC). The resulting systems provided either a detergent-solubilized environment (micelles) or a native membrane-derived lipid environment (nanodiscs) for subsequent experiments. Figure created using BioRender.

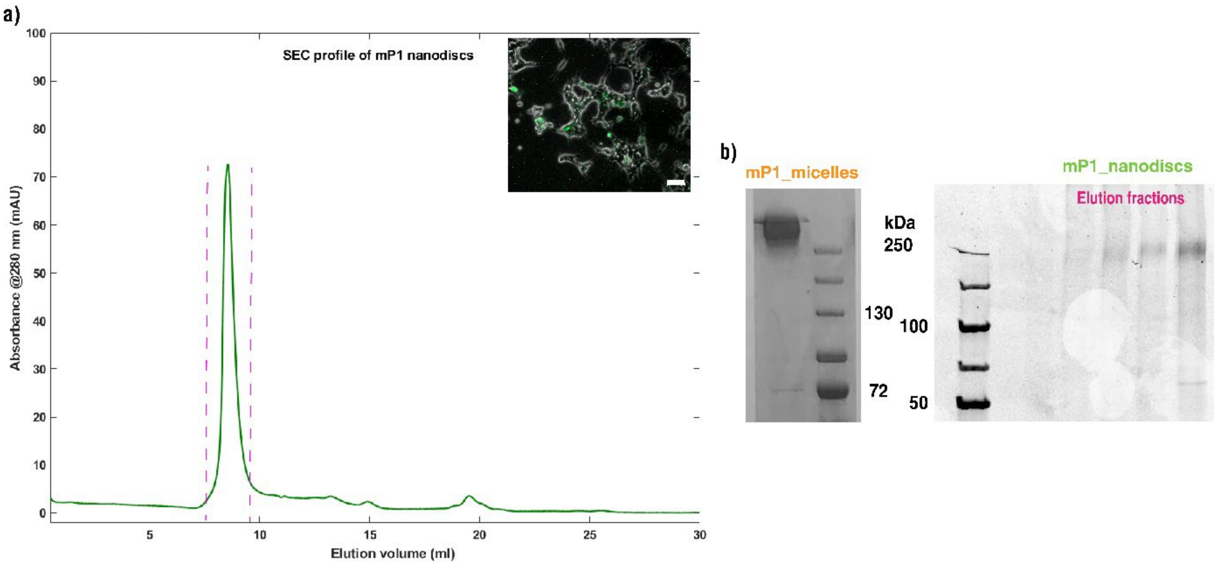


**Fig S2: Characterization of mP1 expression and reconstitution.** (a) Representative size-exclusion chromatography (SEC) profile of purified mP1 nanodiscs. The shaded region indicates the fractions collected for subsequent experiments. Inset: fluorescence microscopy image of HEK293T cells expressing GFP-tagged mP1, shown as an overlay of bright-field and AF488 fluorescence channels. Scale bar: 50 µm. (b) SDS-PAGE analysis of purified mP1 micelles and mP1 nanodiscs. Protein bands migrating above 250 kDa are consistent with the expected molecular weight of full-length PIEZO1. Representative elution fractions obtained following nanodisc purification are shown.

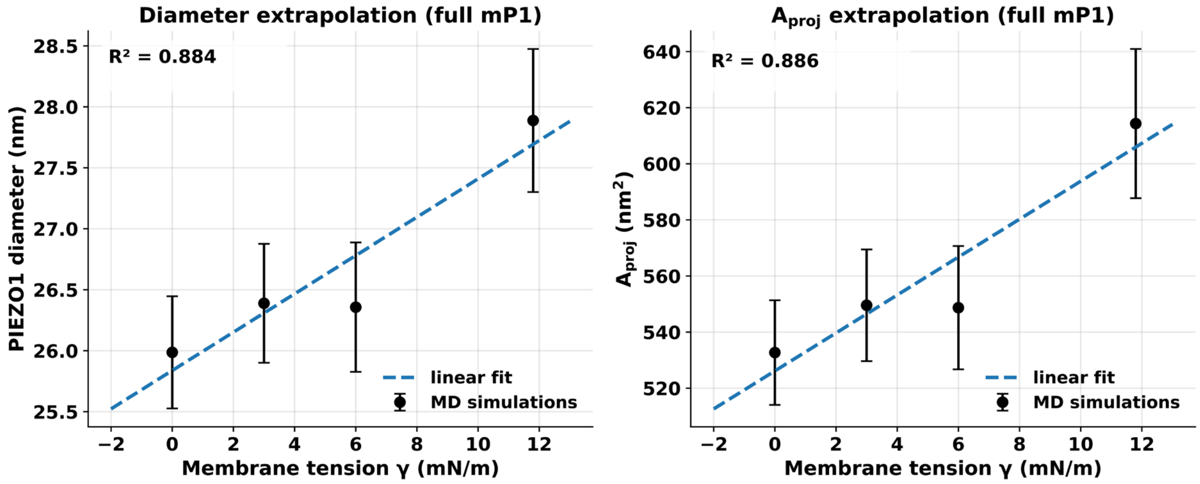


**Fig S3: Estimation of mP1 diameter and projected area under different membrane tensions.** Linear extrapolation of the mP1 diameter (left) and projected area, $A_{proj}$ (right), as a function of membrane tension ($\gamma$). Diameter and area were determined using the full-length mP1 model, in which the center of geometry of the outermost resolved repeat (THU1, residues 13–138) of each monomer was used to define the arm tip (see Methods for details). Data points represent ensemble means ± SEM obtained from ten independent replicas. Dashed lines indicate linear fits used to estimate the corresponding values under compressive membrane tension). Predicted values and coefficients of determination ($R^2$) are shown in each panel.

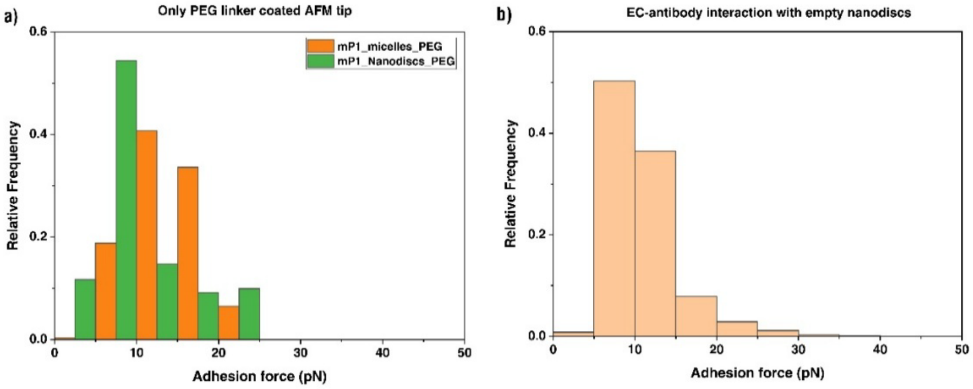


**Fig S4: Control measurements for non-specific interactions in SMFS experiments**. (a) Adhesion-force distributions obtained using AFM tips functionalized with PEG linker only on mP1 micelles and mP1 nanodiscs. (b) Adhesion-force distribution measured using EC-antibody-functionalized AFM tips on empty nanodiscs. In both control experiments, adhesion forces were predominantly below 20 pN, consistent with the noise threshold used for data analysis and substantially lower than the forces measured for specific EC antibody–mP1 interactions.

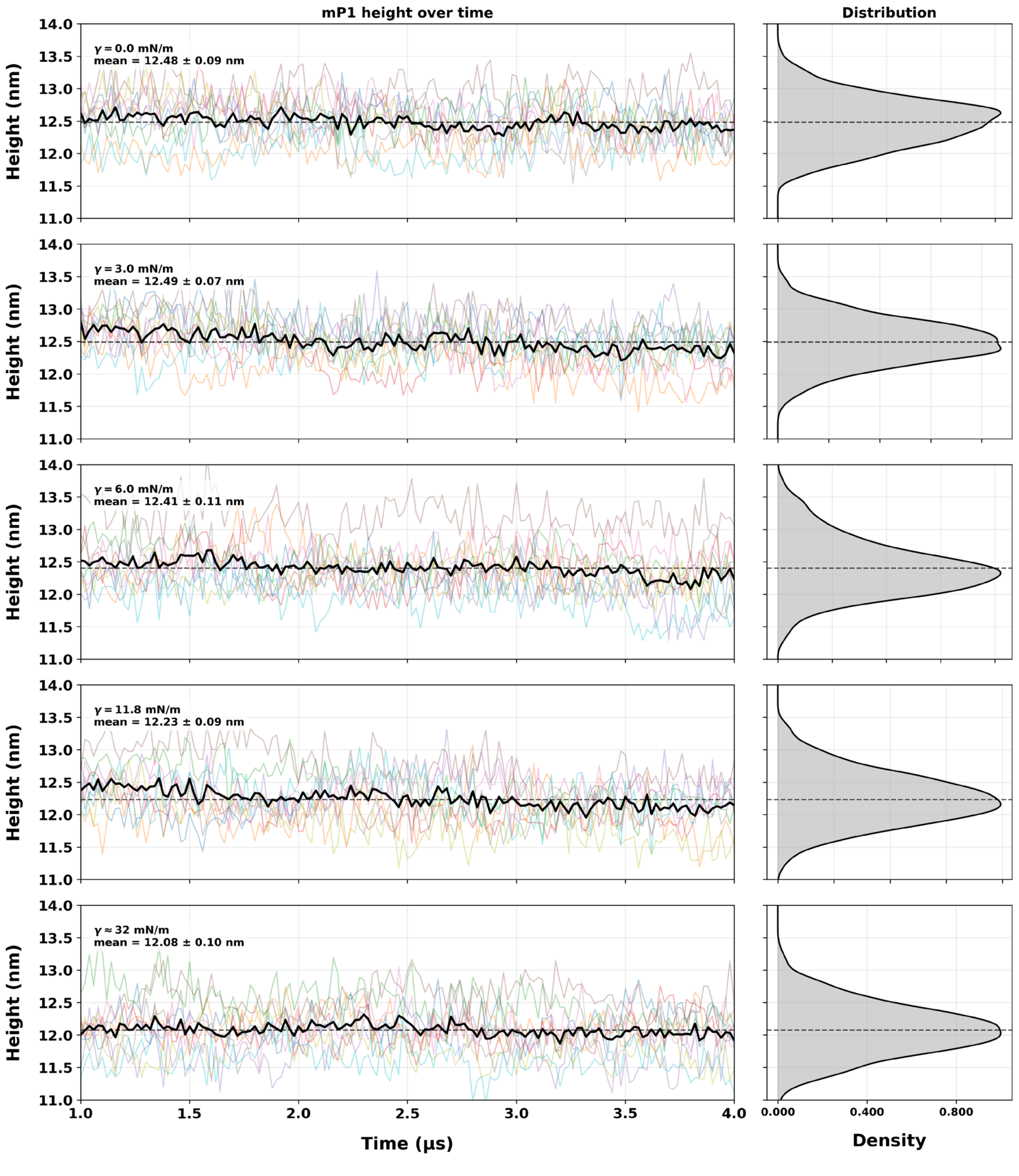


**Fig S5, a: Height analysis of absolute protein under different membrane tensions.** Time series of mP1 total protein height (Cap to CTD) across 10 independent replicas (colored lines) for each membrane tension condition. The black line shows the ensemble mean over time; the dashed line indicates the overall mean. Right panels show the corresponding height distribution. Mean ± SEM values are indicated in each panel.

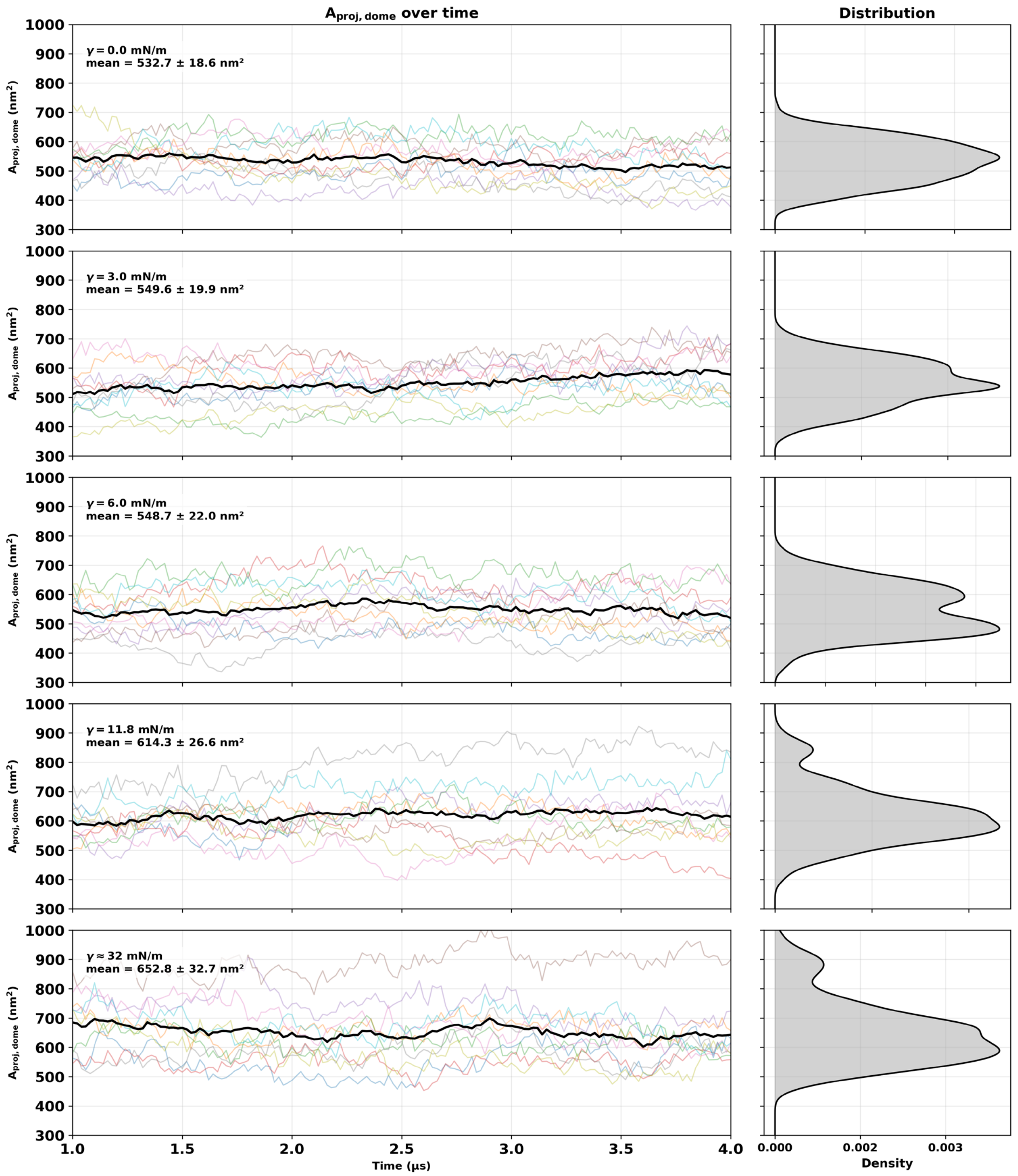


**Fig S5, b: Projected area of mP1 under different membrane tensions (full mP1).** Time series of the mP1 dome projected area (A_proj) across 10 independent replicas (colored lines) for each membrane tension condition. $A_{proj,dome} = \pi \cdot r^2$, where r is the circumscribed radius of the triangle formed by the centre of geometry of the THU1 repeat (residues 13–138) of each monomer chain. The black line shows the ensemble mean over time. Right panels show the corresponding distribution. Mean ± SEM values are indicated in each panel.

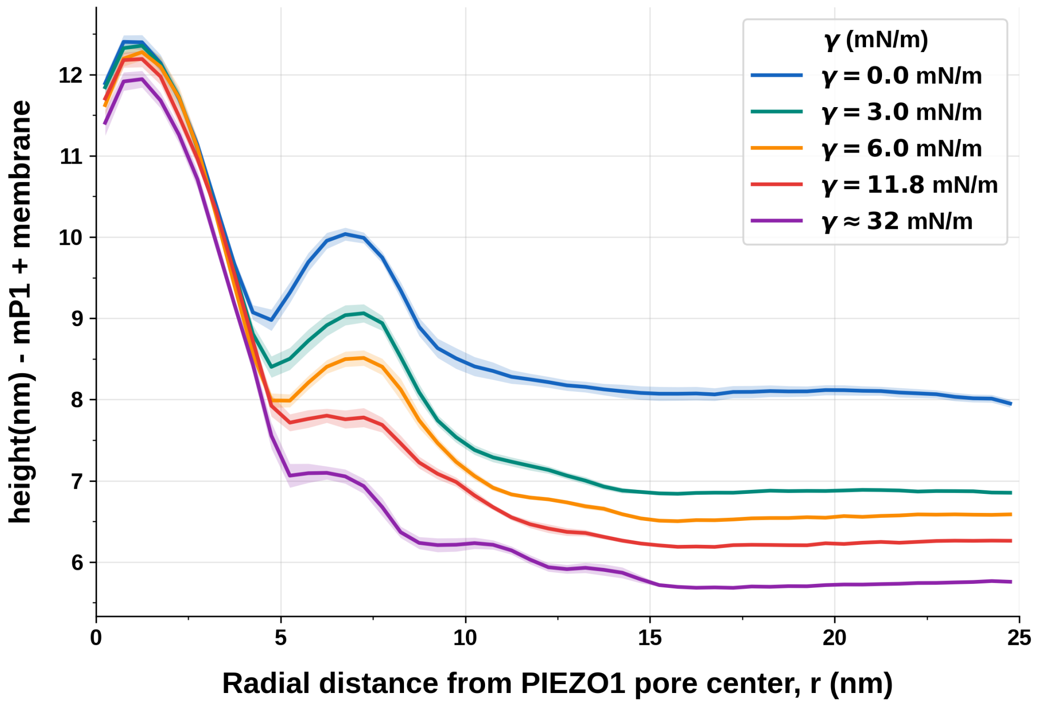


**Fig S6: Radial height profile of mP1 embedded in the lipid bilayer under different membrane tensions.** Mean height of the mP1–membrane system as a function of radial distance from the PIEZO1 pore center, for five membrane tension conditions. Shaded bands indicate ± SEM across 10 independent replicas. The characteristic dome shape at low tension ($\gamma = 0$ mN/m) progressively flattens with increasing membrane tension, consistent with tension-driven conformational changes in the PIEZO1 blade architecture.

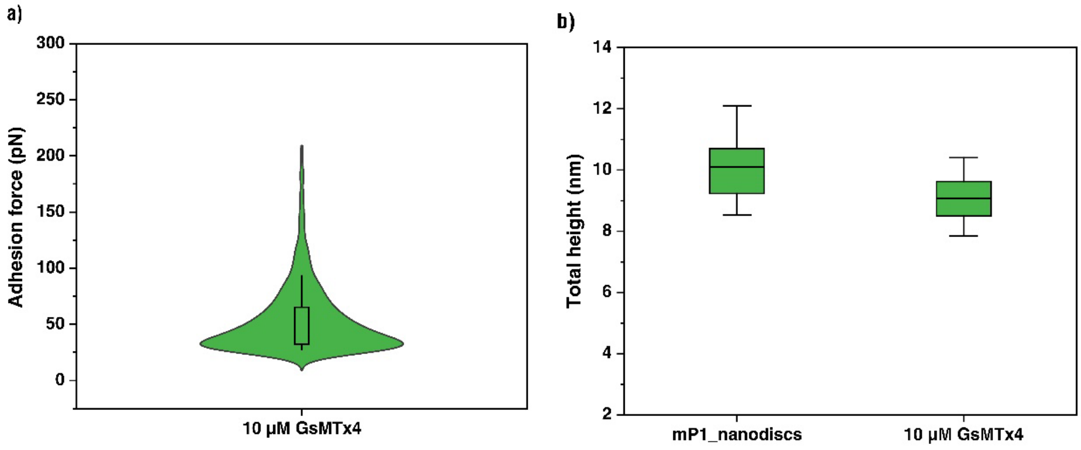


**Fig S7: Effect of GsMTx4 treatment on mP1 nanodiscs.** (a) Adhesion-force distribution measured by SMFS following treatment of mP1 nanodiscs with 10 µM GsMTx4 for 15 min. The distribution exhibited a mean adhesion force of 43.1 ± 3.2 pN. (b) Total particle height of untreated and GsMTx4-treated mP1 nanodiscs measured by AFM. GsMTx4 treatment reduced the mean height from 10.1 ± 1.4 nm in the control to 9.1 ± 0.9 nm, indicating that perturbation of the membrane environment influences the structural and mechanical response of mP1.

**Calculation S1: Estimation of Binding energetics and membrane bending energy**

**a)**

The work done during interaction between PIEZO1 and EC antibody can be approximated to determine the binding energetics. The mechanical work done on the system or the energy utilised can be calculated using

$$E = F.x$$

where E= binding energetics or work on the system

F= forces measured during the protein-antibody interaction i.e. the adhesion force in pN

x=distance measured for the said force (here it is the tip sample separation in nm)

The mean adhesion forces (F) were taken from Fig 3d and Fig 4a. The tip sample separation (x) was approximated from the force curves with representative value shown in Fig 3c.

The average tip-sample separation for mP1 micelles system was ~12 nm and for mP1 nanodiscs was ~14 nm. 1 $k_BT$= 4.1 pN.nm at 298 K was used for thermal energy conversion.

**b)**

The bending energy ($E_b$) for mP1 nanodiscs was calculated using Helfrich model of membrane elasticity[1,2] for flat or slightly curved bilayer.

$$E_b = \frac{K.A}{2}(M - M_0)^2$$

Where $K$= Bending rigidity =35 $k_BT$ for nanodiscs of large diameter[3]

A= area of the nanodisc

M= actual curvature =1/R ; R= calculated mean radii for curvature

$M_0$ = spontaneous curvature =1/$R_0$ ; $R_0$ = spontaneous radii for curvature

For elliptical shaped mP1 nanodiscs as seen through AFM imaging,

Major axis (a) =diameter of the nanodisc=25.88±4.91 nm

Minor axis (b) =12.7±3.2 nm

Area of nanodisc (A)= π*a/2*b/2 ; A= 258.01 $nm^2$; For both leaflets of nanodisc =516.02 $nm^2$

Mean radii of curvature (R) approximated using harmonic mean i.e. $R_{mean}=\frac{ab}{a+b}$ = 8.52 nm

Schachter et al. and Różycki et al estimated that for asymetric bilayer and large nanodiscs, the spontaneous radii range between 10 nm - 35 nm[4-6]. Assuming median value of 22.5 nm for $R_0$

$E_b$ = 48.1 $k_BT$.

For circular nanodiscs (A=525.77 $nm^2$ ; M=0.08 $nm^{-1}$, only one radii i.e. from major axis)

$E_b$ = 11.9 $k_BT$.
